# ‘’Reproductive response of laying chickens to ameliorative method of aflatoxin

**DOI:** 10.1101/2022.06.17.496585

**Authors:** Olayinka Abosede Ojo, Immanuel Bitto, Emmanuel Olubisi Ewuola

**Author notes:** Corresponding author, Department of Animal Production, Fisheries and Aquaculture, Kwara State University.

## Abstract

Aflatoxin is toxic, carcinogenic and ubiquitous in nature, affecting both crops and livestock. Mitigating aflatoxin effect using toxin binders has not been very effective. Information on the use of biological methods in aflatoxin mitigation has not been adequately documented. Consequently, influence of bio-control method of aflatoxin on some reproductive hormones, ovarian weight and histopathological parameters of laying chickens (LC) were investigated. Point-of-lay Bovan Nera (n=700) were harphazardly distributed to four dietary treatments; Aflasafe maize-based diet (AMBD), farm feed (FF), aflatoxin-contaminated diet with toxin binder (ACDTB) and aflatoxin-contaminated diet without toxin binder (ACDWTB). The contaminated diets contained 306.3ppb aflatoxin and the experimental design was completely randomized into four treatments (n= 175) of five replicates (n=35) per treatment for a period of 14 weeks. Blood (5mL) was collected at 14th week for LC to determine the estrogen, luteinising hormone (LH), follicle stimulating hormone (FSH), histopathology of the ovary using standard procedures. Data were analysed using descriptive statistics and ANOVA at a0.05. The ROW (%) ranged from 0.34±0.2 (ACDTB) to 0.93±0.3 (AMBD). Estrogen (mg/dL) value was highest in LC fed ACDWTB (2.75±1.08) and least in FF (2.07±0.52). The LH (iu/L) value was highest in LC fed AMBD (1.29±1.68) and least in ACDTB (0.36±0.32). Histopathology of the ovary showed cysts, observed along the oviduct wall in LC fed ACDTB. AMBD enhanced active laying period in LC with no sign of aflatoxocosis. The use of aflasafe maize grain in poultry diet is recommended.

## 1. Introduction

Aflatoxin (AF) has been confirmed as most toxic, tetratogenic, mutagenic and carcerous mycotoxins so far known (Williams et al., 2009; Khan et al., 2010). The detrimental effect of aflatoxin on various animals has been well documented (Keyl and Booth, 1971; Lawloy and Lynch, 2001; Zain, 2010) and this include yellow ocher discolouration with multifocal haemorrhage with proliferation of the bile ducts (Ehklas, 2012). Hepatomegaly and hypertrophy in some organs and higher relative weights of the organs (Huff et al., 1992; Ologhobo et al., 2015) have been reported. Other effects include accelerated follicular atresia (Siloto et al., 2011), associated by discontinuance of egg laying during the feeding trial (Hafez et al., 1982). Aflatoxin has also been documented to cause disturbances in the hormonal profile of domestic animals, usually resulting in reduced fertility potential (Clarke et al., 1987; Tiemann and Vanselow, 2003; Hasanzadeh et al., 2011). Different approaches have been investigated to mitigate the effect of aflatoxin contamination in livestock feeds. This include use of enteroabsorbants such as hydrated sodium, calcium alumino-silicates. Although, it selectively bind aflatoxin B1 without depleting micronutrients (Williams et al., 2004), but, few successes have recorded with this method. A combination of heat with ammonia has also been stated to irreversibly detoxified aflatoxin (Brown, 2010), but the limitation is that it can only make the blue glow go away, without protecting the feed. Also, microorganisms resembling Mycobacteriumis, Stenotrophomonas maltophilia and Trichoderm viride have also been attested to succeed in breaking down aflatoxins (Liu et al., 2001 and Kasmani et al., 2012). However, they have been noted to pose health risks to consumers. Physical and chemical methods of degradation have been known to cause reduced product nutrient, organoleptic qualities and undesirable health effects (Teniola et al., 2005). Consequent to these limitations, novel bio-control method with biological mechanism has been developed, which involves displacing toxigenic strains of Aspergillus flavus from agricultural fields with strains of Aspergillus flavus that do not produce aflatoxin (atoxigenic strains). This causes encumbrance with the contamination process by physically excluding the toxigenic strain as at the time of microorganism invasion and contending for nutrients needed for aflatoxin biosynthesis by the toxigenic strains, thereby abridging the total toxigenicity of A. flavus population. The significance of this research is to evaluate the changes in reproductive performance during chronic aflatoxicosis and compare it with the use of aflasafe maize-based diet (aflatoxin-free diet) in laying chickens.

## MATERIALS AND METHODS

This study was carried out at God’s Grace Farm, located in Lagun Town, along Ibadan-Iwo road, Oyo State. The Aflatoxin-contaminated maize grain and aflasafe maize grains used for this experiment were obtained from the plant pathology unit, International Institute of Tropical Agriculture, (IITA), Ibadan, Nigeria. Other ingredients used for the feed formulated were obtained from God’s Grace feed Mill, the host farm. Maize grain, used as the aflatoxin carrier was inoculated with toxigenic strain of *Aspergillus flavus* of Nigerian origin. The culturing and inoculation was done at the plant pathology unit, International Institute of Tropical Agriculture (IITA), Ibadan, Nigeria, using 5% V8 juice and 2% agar, with PH 5.2 and a spore load of 2.475 × 106 per ml. Aflatoxins, quantified using scanning densitometer, CAMAG TLC scanner 3 with – CATS 1,4,2 software (Camag AG, Muttenz, Switzerland) (Suhagia et al., 2006)

## EXPERIMENTAL BIRDS AND MANAGEMENT

A total of 700, 30-week old Bovan Nera black hens with a mean body weight of 2.0kg was used for this experiment The experimental birds were randomly allotted into 4 treatments consisting of 175 birds per treatment, replicated thrice with 35 birds/ replicate. The laying chickens were housed in battery cages having linear feed troughs and nipple drinkers for running water. The birds were fed basal feed for 14 days (acclimatization period), after which experimental diets and fresh water were provided *ad libitum*. Data collection was done weekly throughout the duration of the study which lasted for 14 weeks

### 2.2 EXPERIMENTAL DIETS

Four (4) experimental diets were formulated based on nutrient requirement of laying hens. The experimental diets are as shown in Table 1 comprising of the following; Treatment 1 (AMBD) – Aflasafe maize-based diet, Treatment 2 (FF+Toxin binder) –Farm feed with toxin binders (Control diet), Treatment 3 ((ACDTB)– Aflatoxin contaminated diet with toxin binder, Treatment 4 (ACDWTB)– Aflatoxin-contaminated diet without toxin binder. The diets were subjected to chemical analysis, to obtain proximate composition as stated in Table 1 under the Analyzed nutrient segment. At the end of the experiment (14th week), 80 birds were randomly selected from each treatment and three (3) mls of blood sample was collected through the jugular vein in the earliest hours of the day, into anticoagulant-free bottles. These bottles were kept in slanting position and allowed to clot. The samples were spun at 3000 rpm for 10 minutes, serum samples obtained were then separated into sterile tubes for analysis. Enzyme Linked Immunosorbent Assay (ELIZA) kits were used for the assay of estrogen, follicle stimulating hormone and luteinizing hormone. Equipment used was ELIZA microplate reader/ absorbance (IRE 96), a fully automated 8-channels measurement system having absorbance as its detection mode (Bourne et al., 2003). After feeding trial, 80 birds (20 birds per replicate) were randomly selected per treatment, weighed and sacrificed. Dissection was done through the lower abdominal incision. Samples of the ovary and reproductive organ were harvested for histopathological investigation. The tissues were cut into small pieces of not more than 4mm thick and placed into pre-labelled cassettes. These were further immersed in 10% formal saline for 24 hours to fix. Tissue processing was done using hematoxylin and eosin technique (Avwioro, 2010).

**Table 1:**
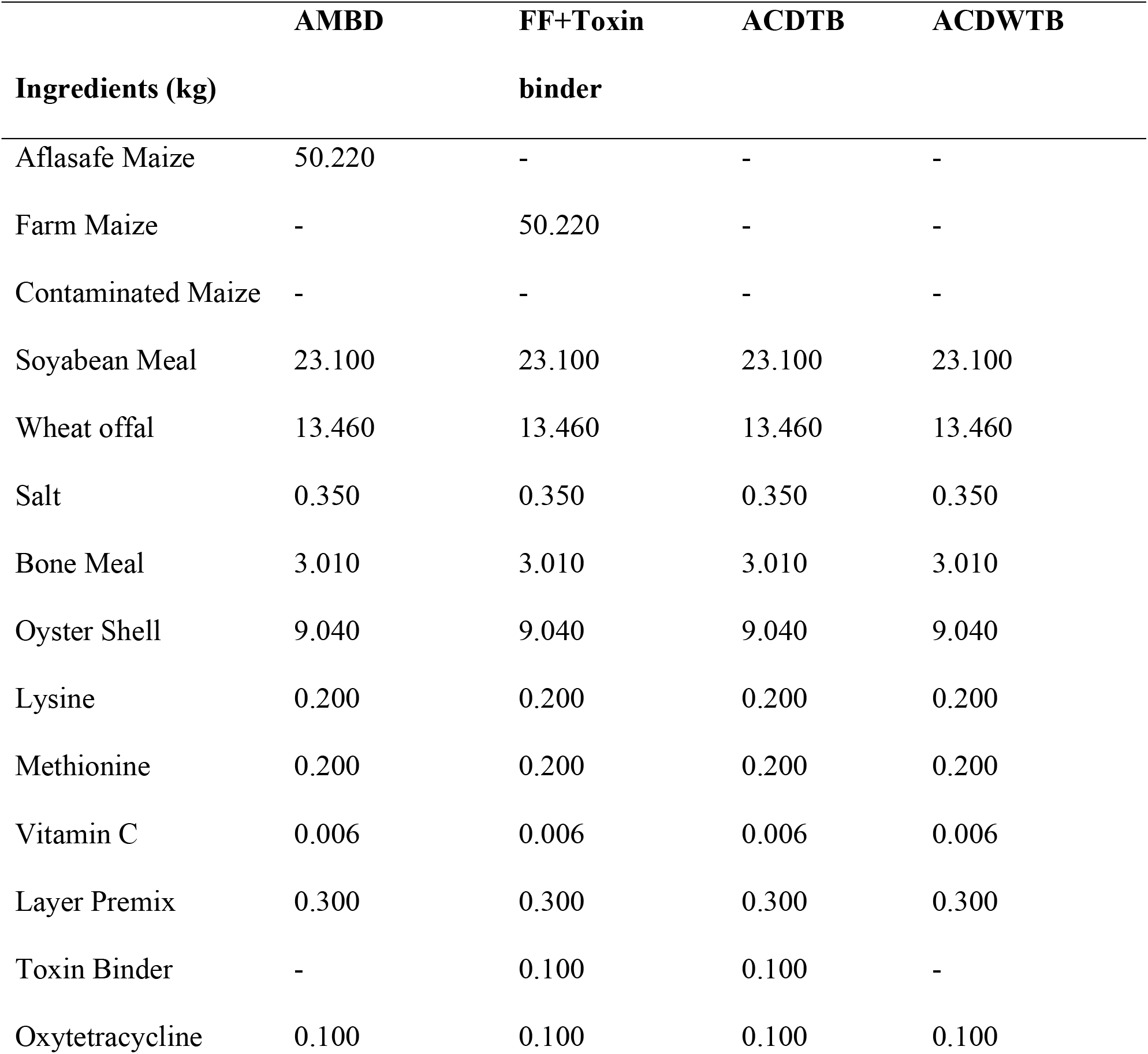
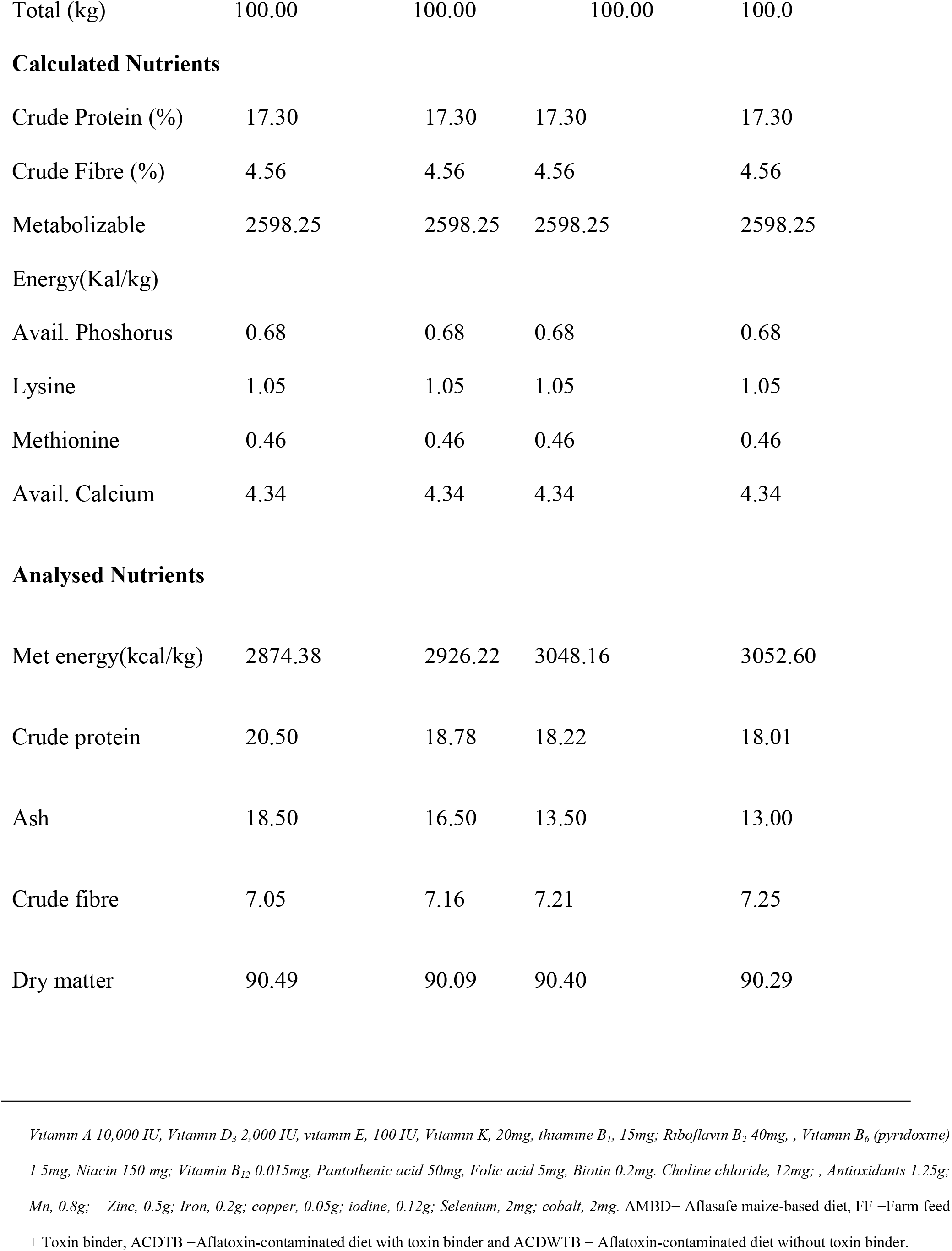
Composition of the experimental diet (%) (Layers Mash)

|  | **AMBD** | **FF+Toxin** | **ACDTB** | **ACDWTB** |
| --- | --- | --- | --- | --- |
| **Ingredients (kg)** |  | **binder** |  |  |
| Aflasafe Maize | 50.220 | - | - | - |
| Farm Maize | - | 50.220 | - | - |
| Contaminated Maize | - | - | - | - |
| Soyabean Meal | 23.100 | 23.100 | 23.100 | 23.100 |
| Wheat offal | 13.460 | 13.460 | 13.460 | 13.460 |
| Salt | 0.350 | 0.350 | 0.350 | 0.350 |
| Bone Meal | 3.010 | 3.010 | 3.010 | 3.010 |
| Oyster Shell | 9.040 | 9.040 | 9.040 | 9.040 |
| Lysine | 0.200 | 0.200 | 0.200 | 0.200 |
| Methionine | 0.200 | 0.200 | 0.200 | 0.200 |
| Vitamin C | 0.006 | 0.006 | 0.006 | 0.006 |
| Layer Premix | 0.300 | 0.300 | 0.300 | 0.300 |
| Toxin Binder | - | 0.100 | 0.100 | - |
| Oxytetracycline | 0.100 | 0.100 | 0.100 | 0.100 |
| Total (kg) | 100.00 | 100.00 | 100.00 | 100.0 |
| **Calculated Nutrients** |  |  |  |  |
| Crude Protein (%) | 17.30 | 17.30 | 17.30 | 17.30 |
| Crude Fibre (%) | 4.56 | 4.56 | 4.56 | 4.56 |
| Metabolizable | 2598.25 | 2598.25 | 2598.25 | 2598.25 |
| Energy(Kal/kg) |  |  |  |  |
| Avail. Phosphorus | 0.68 | 0.68 | 0.68 | 0.68 |
| Lysine | 1.05 | 1.05 | 1.05 | 1.05 |
| Methionine | 0.46 | 0.46 | 0.46 | 0.46 |
| Avail. Calcium | 4.34 | 4.34 | 4.34 | 4.34 |
| **Analysed Nutrients** |  |  |  |  |
| Met energy(kcal/kg) | 2874.38 | 2926.22 | 3048.16 | 3052.60 |
| Crude protein | 20.50 | 18.78 | 18.22 | 18.01 |
| Ash | 18.50 | 16.50 | 13.50 | 13.00 |
| Crude fibre | 7.05 | 7.16 | 7.21 | 7.25 |
| Dry matter | 90.49 | 90.09 | 90.40 | 90.29 |

*Vitamin A 10,000 IU, Vitamin D<sub>3</sub> 2,000 IU, vitamin E, 100 IU, Vitamin K, 20mg, thiamine B<sub>1</sub>, 15mg; Riboflavin B<sub>2</sub> 40mg, , Vitamin B<sub>6</sub> (pyridoxine) 1 5mg, Niacin 150 mg; Vitamin B<sub>12</sub> 0.015mg, Pantothenic acid 50mg, Folic acid 5mg, Biotin 0.2mg. Choline chloride, 12mg; , Antioxidants 1.25g; Mn, 0.8g; Zinc, 0.5g; Iron, 0.2g; copper, 0.05g; iodine, 0.12g; Selenium, 2mg; cobalt, 2mg. AMBD= Aflasafe maize-based diet, FF =Farm feed + Toxin binder, ACDTB =Aflatoxin-contaminated diet with toxin binder and ACDWTB = Aflatoxin-contaminated diet without toxin binder.*

**Table 2:**
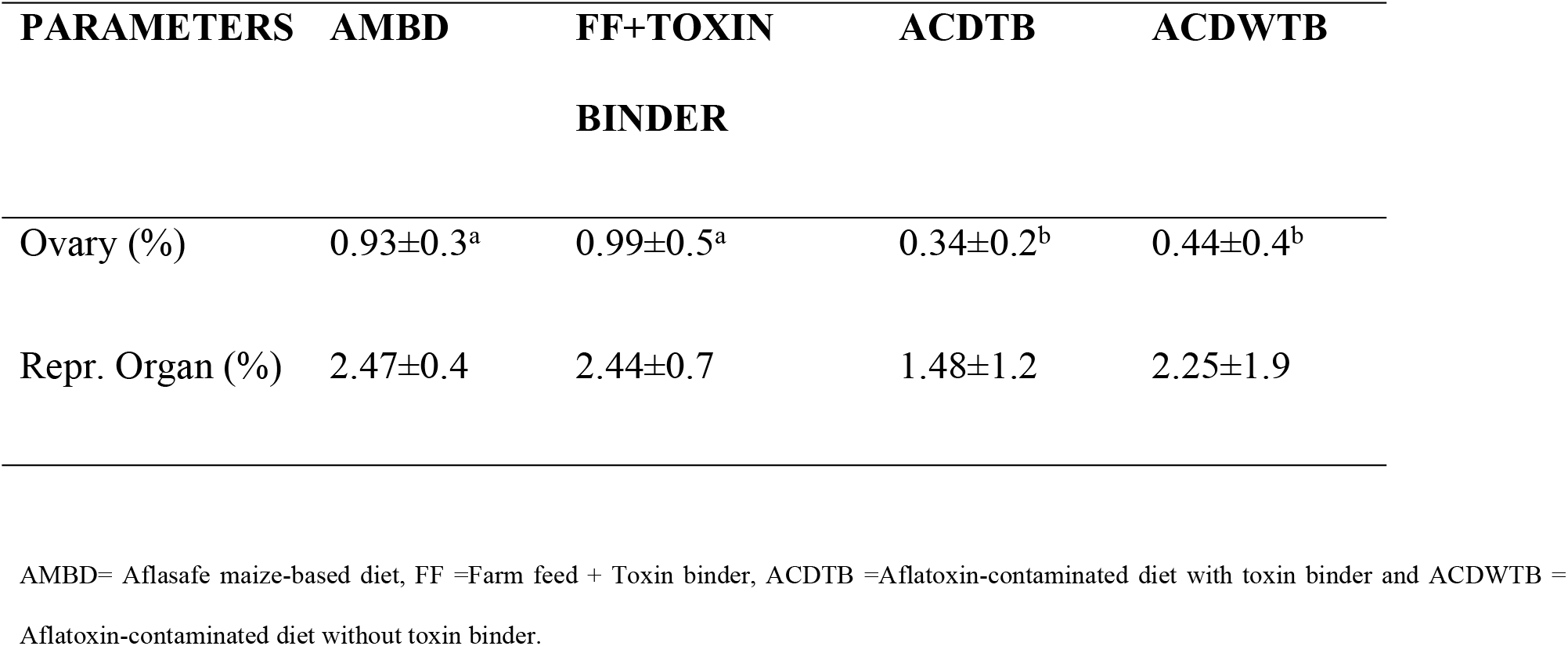
Weight of Reproductive Organ of laying chickens fed experimental diets.

### 2.3 Statistical Analysis

All data obtained from the study was subjected to descriptive statistics and one-way analysis of variance (ANOVA) in a completely randomized design, using statistical analysis software (SAS, 2008). Means were separated using Duncan multiple range test (Duncan, 1957).

## 4. Results and discussion

The relative ovary weight of birds fed AMBD (0.93%)) and FF+toxin binder (0.99%) were not different, however, both were significantly higher than the values observed in birds which received ACDTB (0.34%) and ACDWTB (0.44%) respectively. The result is in tandem with the observation of Ajani et al. (2014), who noticed ovary atresia i.e degeneration of immature ovarian follicles during follicular phase in laying hens fed a diet containing 8000ppb AFB for 7 days. According to Hafez et al. (1982), aflatoxicosis causes pathological changes in the chicken ovaries, which has detrimental effect on egg production. Therefore, it can be inferred that aflatoxin could have adverse effect on egg production, due to aflatoxin-related pathological changes in the ovary as reported in this study. The chickens fed ACDTB, showed ovarian cycts along the oviduct wall. This could also have been due to the aflatoxin-contamination effect. Cysts are fluid-filled cavities that may occur along the oviduct wall, resulting into tumor, when larger than 2mm in diameter. It has been noted that production of progesterone is usually by the largest growing follicles and its receptors are found along different points in the oviduct wall. The observed cycts could have blocked or reduced the number of progesterone receptors, causing a depression effect in the mechanism of progesterone production and its activity, with a negative influence on oviduct contraction and egg transport. Perhaps, this have caused a drastic reduction in egg oviposition, resulting into re-absorption of yolk into the system of the hens. This result is in accordance with the reports of El-Azab et al. (2009) and Hasanzadeh et al. (2011), who stated that, aflatoxin has been shown to disrupt the reproductive system in both male and female animals, causing alterations in the form of growing and mature ovarian follicles.

### 4.1 Dietary effect on hormonal assay of laying chickens

The results of the luteinizing hormone (LH), estrogen and follicle stimulating hormone (FSH) assays of laying chickens fed experimental diets during treatment period are illustrated in Table 3. Except the FSH values that was not significant, the luteinizing and estrogen hormone values showed significant differences. The LH value (1.29±1.68 IU/L) obtained in birds fed the AMBD, was not significantly (P<0.05) different compared with those given FF+toxin binder (0.64±1.27 IU/L), the LH values (0.36±0.32 IU/L and 0.42±1.03 IU/L) of laying birds fed aflatoxin-contaminated diets with and without toxin binder were significantly lower. The mean value of estrogen recorded in birds fed AMBD (2.10±0.52pg/mL) showed no influence compared to the FF+toxin binder (2.07±0.83pg/mL). The mean values recorded for birds fed ACDTB (2.38±0.90pg/mL) and ACDWTB (2.75±1.08pg/mL) were not statistically influenced by the dietary treatment. The FSH values of laying chicken were not significantly (P<0.05) different among the treatments.

**Table 3:** Hormonal Assay of laying hens fed experimental diets.

| PARAMETERS | T1 | T2 | T3 | T4 |
| --- | --- | --- | --- | --- |
| Luteinizing hormone (LH) | $1.29 \pm 1.68^a$ | $0.64 \pm 1.27^{ab}$ | $0.36 \pm 0.32^b$ | $0.42 \pm 1.03^b$ |
| Estrogen | $2.10 \pm 0.52^b$ | $2.07 \pm 0.83^b$ | $2.38 \pm 0.90^{ab}$ | $2.75 \pm 1.08^a$ |
| Follicle Stimulating | 2.66±1.33 | 2.24±0.84 | 2.13±0.68 | 2.22±0.79 |
| Hormone (FSH) |  |  |  |  |

ab: means on the same rows but with different superscripts are significant ( $P < 0.05$ ). AMBD = Aflasafe maize-based diet, FF+Toxin binder = Farm feed + toxin binder, ACDTB = Aflatoxin-contaminated diet + toxin binder, ACDWTB = Aflatoxin-contaminated diet without toxin binder.

**Table 4:** Histopathological examination of the ovary given aflatoxin-contaminated diet.

| Parameter | AMBD | FF+Toxin<br>binder | ACDTB | ACDWTB |
| --- | --- | --- | --- | --- |
| Ovary | No visible<br>lesion | No visible<br>lesion | Cyst observed<br>along the<br>oviduct wall | Cyst observed<br>along the<br>oviduct wall |

Reproductive hormones of birds fed experimental diet was significantly influenced by the dietary treatment. Luteinizing hormone is a primary hormone of reproduction, produced from the anterior lobe of the pituitary gland. The epithelial cells lining the empty follicular cavity has been reported to multiply under the influence of luteinizing hormone, to form corpus luteum. In poultry, ovulation and oviposition processes are controlled by luteinizing hormone and progesterone. A higher concentration of serum LH of experimental laying birds, as observed in birds fed AMBD, is an advantage as it would improve the egg laying capacity of the birds which would translate into higher profitability. The depressed values of LH in this study corroborate the findings of Hasanzadeh et al. (2011) and Ismail (2012), who reported a significantly lowered luteinizing hormone level in the aflatoxin-treated groups of male rats compared to the control, which received zero dosage of aflatoxin. This result might be attributed to the adverse effect of AF on the hypophysis, resulting into hypophysotoxicity, according to Clarke et al. (1987). Estrogen is a steroid hormone produced in the gonads, which performs a prominent role in the development of female secondary sex characteristics and in ovarian cycle. Trophic hormones FSH and LH are noted for stimulating the synthesis and release of estradiol from the theca interna of matured follicle. It has been documented that toxins cause a cascade of events called acute phase response, which induced local inflammation and activation of the hypothalamic-pituitary-adrenal axis (HPA), as stated by Sheng *et al*. (2021). Perhaps, the local inflammation of the HPA caused by AF effect had resulted into a significantly lowered concentration of LH and FSH in laying birds, which could have caused a concormitant decrease in the estrogen value. Documentation showed that FSH and LH release from the pituitary gland occurs in a rhythmic manner, as regulated by the hypothalamic biologic clock, known as pulse frequency (Squires, 2003). The frequency of GnRH (Gonadotropic Releasing Hormone) by the hypothalamus, differentially affect FSH and LH release, which should also influence estrogen synthesis and release. Contrary to this, the estrogen level of birds in ACDWTB is significantly higher compared to the control (FF+Toxin binder). In literature, it has been stated that cholesterol is the precursor of estradiol and it is present as low density lipoprotein (LDL) in plasma. In the serum biochemical response of hens, the cholesterol level of the birds in ACDWTB is highest. Perhaps, the high level of LDL (low density lipoprotein) in the plasma of the laying hens acted as a precursor for increased synthesis of estrogen in the laying birds in ACDWTB, with the recorded highest value. FSH functions in maintaining the hierarchy of size in the developing follicles and the rate of follicular atresia. It also works synergistically with luteinizing hormone to cause an elaborate secretion of estrogen during ovulation. An increased FSH concentration in birds fed AMBD suggested a normal hierarchical follicular development and ovulation, as well as normal egg production (Squires, 2003).

**Plate 1:**
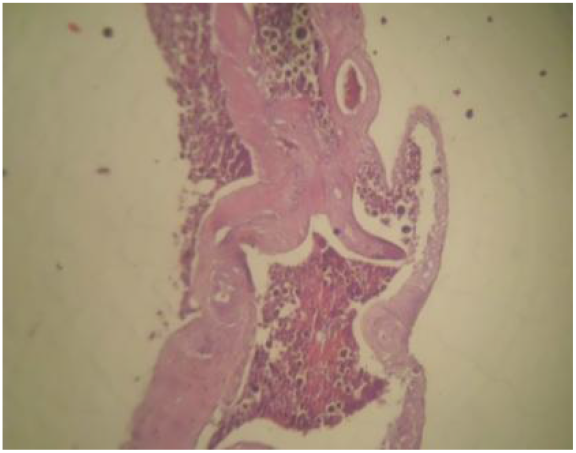
Showing the Ovary of laying chickens fed AMBD. No visible lesion observed.

**Plate 2:**
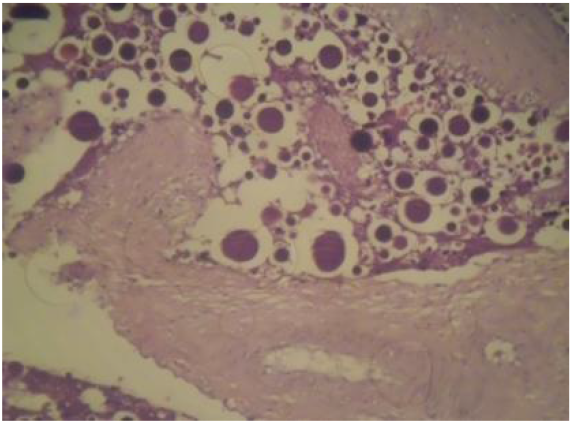
Showing the Ovary of laying chickens fed FF. No visible lesion was observed.

**Plate 3:**
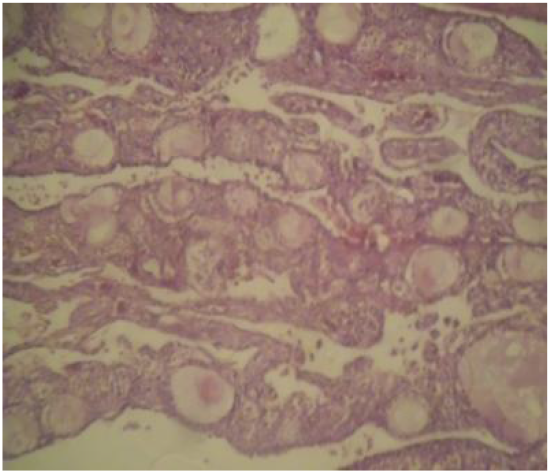
Showing the Ovary of laying chickens fed AF-CDTB. No visible lesions seen. There are cysts along the oviduct wall.

**Plate 4:**
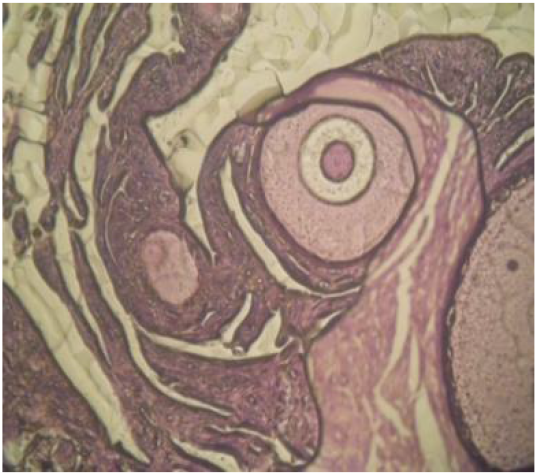
Showing the ovary of laying chicken fed AF-CDTWB. There is no observable visible lesion.

### 4.2 Ovary Histopathology

The microscopic investigation of various organs in previous studies showed that aflatoxin adversely affected the organs attributed with the hematopoietic, immune and the reticuloendothelial system (Qureshi et al., 1998; Ortatatli and Oguz, 2001; Ortatatli et al., 2005 and Eklas, 2012). The ovary of laying chickens fed AMBD, which showed no histopathological alterations may be due to the aflatoxin-free diet obtained through the use of aflasafe maize. Similar result was observed in birds fed the FF+Toxin binder diet and birds fed ACDTWB. Although the ACDTWB diet had a recorded high aflatoxin concentration, which resulted into cessation of egg production in the group, corroborating the findings of Hafez et al. (2008) and Ismail (2012) who observed that aflatoxin-treated mature domestic fowls showed follicular atresia, accompanied by cessation of egg production during the whole feeding period. Hasanzadeh et al. (2014) also observed that the micromorphological effects of the AFB1 on the ovary showed that the attic changes were seen in different ovarian follicular layers, including oocyte, granulose and theca, which increased both at the microscopic and macroscopic level (Siloto et al., 2011). Yet no visible lesion was observed and the concentration of the aflatoxin used showed no histopathological alteration on the ovary. Aflatoxin has also been shown to disrupt the reproductive system in both male and female animals, causing alterations in the form of the growing and mature ovarian follicles. (El-Azab et al., 2009; Hasanzadeh et al., 2014).

## Conclusion and recommendation

Adverse effects of aflatoxin in laying chickens can be prevented by the use of feed ingredients that contains no aflatoxin contamination through the mutual exclusion mechanism of *Aspergillus species* in the soil. Although aflatoxin binder is effective, but its cost implication could increase the overall production cost, especially on a large scale livestock farming. Aflatoxicosis in livestock can be mitigated with the use of aflasafe maize-based diets, it is therefore recommended for use in both small and large scale production.

## Ethics Statement

This research was carried out in strict accordance with the recommendations in the Guide for the Care and Use of Laboratory Animals of the National Institutes of Health. The protocol was approved by the Committee on the Ethics of Animal Experiments of the Kwara State University.

## Consent for publication

“Not applicable”

## Availability of data and materials

The data sets used and/or analysed during the current study are available from the corresponding author on reasonable request.

## Authors’ contributions

**IB** made immense contributions to the design of the study and drafted the hormonal inclusion in this research; **EO** made the collaborative funding acquisition for this research possible and substantively revised it till perfection,**OA** collected, statistically analyzed and interpreted the data used in writing this manuscript and was a major contributor in writing this manuscript.

## Acknowledgements

The authors of this manuscript gracefully acknowledged the expertise and advice that all the contributors have given to developing this study. These include; All Staff of the Pathological Unit, International Institute of Tropical Agriculture (IITA) and all staff and management of God’s Grace Farm, Lagun, Oyo State, Nigeria.

## REFERENCES

Ajani Sudheer, D. V., Tanuja, P. and Pasha K. U. (2014). Aflatoxins. Indian Journal of Advances in chemical Science 3 (2014) 49–60. https://www.ijacskros.com

Avwioro O. G. (2010). Histochemistry and tissue pathology, principle and techniques, Claverianum press, Nigeria.

Brown D. Food chain Mycotoxins 2010: Threats and solutions.

Bourne, M., Franco, M., Wilkes, J. (2003), “Corporate performance measure-ment”, Measuring Business Excellence, Vol. 7 No.3, pp. 15–21.

Clark JD, Jain AV, Mahaffey EA. Experimentally induced chronic aflatoxicosis in rabbits. American Journal of Veterinary Research, 1980 41: 1841–1845.

Clarke RN, Doerr JA, Ottinger, MA.. Age-Related Changes in Testicular Development and Reproductive Endocrinology Associated with Aflatoxicosis in the Male Chicken1 Biology of Reproduction, 1987 Volume 36, Issue 1, 1 February 1987, Pages 117–124, https://doi.org/10.1095/biolreprod36.1.117

Duncan, O. D. The Negro Population of Chicago: A Study of Res-idential Succession, Chicago: University of Chicago Press.1977, Introduction to Structural Equation Models, New Yor: Academic press.

Ekhlas KH. Histopathological changes of some internal organs in broilers fed aflatoxin. Al-Qadisiya Journal of Vet. Med. Sci. 2012 Vol./11.No./2 2012.

El-Azab SM. Study of aflatoxin B1 as a risk factor that impair the reproductive performance in females-Egypt. The Internet Journal of Toxicology, 2009 http://264Aflatoxins-RecentAdvancesandFutureProspects. https://www.ispub.com:80/jourbal/the-internet-journal-of-toxicology/1.

Hafez AH, Megalla SE, Abdel-Fattah HM, Kamel YY.: Aflatoxin and aflatoxicosis II. Effects of aflatoxin on ovaries and testicles in mature domestic fowls. Mycopathologia, 1982 77:137–139.

Hasanzadeh SH, Hosseini E. Rezazadeh L. Effects of aflatoxin B1 On profiles of gonadotropic (FSH and LH), steroid (testosterone and 17ß;-estradiol) and prolactin hormones in adult male rat. Iran Journal of Veterinary Research, Shiraz University, 2011 vol. 12, No. 4, Ser. No. 37, 2011.

Huff W. E., Kubena L. F., Harvey R.B., Philips T.D. (1992). Efficacy of hydrated sodium calcium aluminosilicate to reduce the individual and combined toxicity of aflatoxin and ochratoxin A. Poult Sci 71: 64–69.

Ismail NH. Assesment of DNA damage in testes from young Wistar male rats treated with monosodium glutamate. Life Science Journal 2012 9, 930–939.

Kasmani FB, Krimi MAT, Allameh, A. Shariatmadari F. A novel aflatoxin-binding Bacillus probiotic: Performance, serum biochemistry and immunological parameters in Japanese quail. Poultry Science. 2012 91 (8): 1846–1853. Doi:10.3382/ps.2011-01830.

Keyl AC, Booth, AN. Aflatoxin effects in livestock. J. Amer. Oil Chem. Soc. 1971 48: 599–604.

Khan WA, Khan MZ, Khan A, Hussain I. Pathological effects of aflatoxin and their amelioration by vitamin E in White Leghorn layers. Pak. Vet. J. 2010 30 (3):155–162.

Lawloy PG, Lynch. Mycotoxins in pig feed 1: Source of toxins, prevention and management of mycotoxicosis. Irish Vet. J. 2001 54: 67–71.

Liu, J., C. Yang, S. Wasser, H. Shen, C. Tan and C. Ong, 2001. Protection of salvia miltiorrhiza against aflatoxin-B1-induced hepatocarcinogenesis in Fischer 344 rats dual mechanisms. Life Sci., 69: 309–326. Direct Link |

Ologhobo AD, Ewuola EO, Jerome UU, Franca UO., Ifarajimi Osaa. Growth, Nutrient Digestibility of Broilers fed Aflatoxin Contaminated Diets with Aflatoxin Binders. ARPN Journal of Science and Technology.Vol. 5, No. 5, May 2015.ISSN 2225-7217.2

Ortatatli M, Oguz H. Ameliorative effects of dietary dipnoptilolite on pathological changes in broiler chickens during aflatoxicosis. Res. Vet. Sci. 2001 71: 59–66.

Ortatatli M, Oguz H, Hatipoglu F, Karaman M. Evaluation of pathological changes in broilers during chronic aflatoxin (50 and 100 ppb) and clinoptilolite exposure. Research in Veterinary Science, 2005 78, 61–68.

Siloto E V, Sartori D R S, Olivera E F A, Sartori J R, Fascina V B and Berto D A. 2011. Performance and Egg Quality of Laying Hens Fed Diets Containing Aflatoxin, Fumonisin and Adsorbent. Brazilian Journal of Poultry Science 13(1) 21–28.

Qureshi MA, Brake J, Hamilton PB, Hagler WM. Nesheim S. Dietary exposure of broiler breeders to aflatoxin results in immune dysfunction in progeny chicks. Poult. Sci. 1998 77:812–819.

Sheng JA. Bales NJ, Myers SA, Bautista A.I, Roueinfar M, Hale T.M and Handa RJ.. The Hypothalamic-Pituitary-Adrenal Axis: Development, Programming Actions of Hormones, and Maternal-Fetal Interactions Front. Behav. Neurosci., 13 January 2021 https://doi.org/10.3389/fnbeh.2020.601939

Siloto EV, Sartori DRS., Olivera EFA, Sartori JR, Fascina VB, Berto DA. Performance and Egg Quality of Laying Hens Fed Diets Containing Aflatoxin, Fumonisin and Adsorbent. Brazilian Journal of Poultry Science 2011 13(1) 21–28.

Suhagia BN, Shah SA, Rathod IS, Patel HM, Shah, DR, Marolia, BP. Determination of gatifloxacin and ornidazole in tablet dosage forms by high-performance thin-layer chromatography. Anal Sci. 2006: 22 (5):743–745. doi: 10.2116/analsci.22.743.

Squires EJ. Applied Animal Endocrinology. CAB International, 2003 Cambridge, M.A 02139, USA.

Teniola, O. D., Addo, P. A., Brost, I. M., Farber, P., Jany, K. D., Alberts, J. F., et al. (2005). Degradation of aflatoxin B1 by cell-free extracts of Rhodococcus erythropolis and Mycobacterium fluoranthenivorans sp. nov. DSM44556T. Int. J. Food Microbiol. 105, 111–117. doi: 10.1016/j.ijfoodmicro.2005.05.004 PubMed Abstract | CrossRef Full Text | Google Scholar

Tiemann US, Vanselow J. Effect of the mycotoxin and beta zearalenol on regulation of progesterone synthesis in cultured granulose cells from porcine ovaries. Reproductive Toxicology, 2003. 17, 6:673-681.

Williams JH, Philips TD, Jolly CM, Aggarwal D. Human aflatoxicosis in developing countries: a review of toxicology, exposure, potential health consequences and interventions. American journal of Clinical Nutrition 2004 80, 1106–1122.

Williams DE, Orner G, Williard KD, Tilton S, Hendricks JD. Ranibow trout (Oncorhynchus mykiss) and ultra-low dose cancer studies. Comp.Biochem. Physiol. C, 2009, 149: 175–181. PMID : 19135172

Zain, ME.. Impact of mycotoxins on humans and animals. Original Article. Journal of Saudi Chemical Society 2010 15, 129–144.

